# Transient Compartmentalization dynamics in the presence of mutations and noise

**DOI:** 10.1101/521211

**Authors:** Alex Blokhuis, Philippe Nghe, Luca Peliti, David Lacoste

**Author notes:** Corresponding author Email address (Alex Blokhuis).

## Abstract

We extend a recently introduced framework for transient compartmentalization of replicators with selection dynamics, by including the effect of mutations and noise in such systems. In the presence of mutations, functional replicators (ribozymes) are turned into non-functional ones (parasites). We evaluate the phase diagram of a system undergoing transient compartmentalization with selection. The system can exhibit either coexistence of ribozymes and parasites, or a pure parasite phase. If the mutation rate exceeds a certain level called the error threshold, the only stable phase is the pure parasite one. Transient compartmentalization with selection can relax this error treshold with respect to a bulk quasispecies case, and even allow ribozymes to coexist with faster growing parasites.

In order to analyze the role of noise, we also introduce a model for the replication of a template by an enzyme. This model admits two regimes: a diffusion limited regime which generates a high noise, and a replication limited regime, which generates a low noise at the population level. Based on this model, we find that, since the ribozyme dynamics belongs to the replication limited regime, the effects of noise on the phase diagram of the system are mostly negligible. Our results underlines the importance of transient compartmentalization for prebiotic scenarios, and may have implications for directed evolution experiments.

## 1. Introduction

Compartments play a central role in many biological processes of cells, in particular in organelles such as the ER or in the Golgi apparatus [1]. Cells use compartments to organize chemical reactions in space: compartments eliminate the risk of losing costly catalysts which are essential for biochemical reactions, they also accelerate chemical reactions, while reducing the risk of cross-talks due to other side reactions.

In the early 20th century, Oparin suggested that membrane-less compartments, which he called coacervates, could have played a central role in the origin of life [2]. Many compartments in cells are bounded by membranes, but following the recent discovery of the so-called P-granules in C elegans embryo [3], biologists noticed that membrane-less compartments abound in living organisms. These membrane-less compartments are of interest for physicists who want to understand better the active non-equilibrium phase separation which creates them [4]; but also for chemists who are trying to synthetize and control them *in vitro* [5, 6].

After the discovery of the structure of DNA, the coacervates scenario for the origin of life got less popular, and was replaced by replication scenarios [7, 8]. In the sixties, Spiegelman showed that RNA could be replicated by an enzyme called *Qβ* RNA replicase, in the presence of free nucleotides and salt [9]. After a series of serial transfers, he observed the appearance of shorter RNA polymers, which he called parasites. Typically, these parasites are non-functional molecules which replicate faster than the RNA polymers introduced at the beginning of the experiment. In 1971, Eigen conceptualized this observation by proving theoretically that for a given accuracy of replication and a relative fitness of parasites, there is a maximal genome length that can be maintained without errors [10]. This result led to the following paradox: to be a functional replicator, a molecule must be long enough. However, if it is long, it cannot be maintained since it will quickly be overtaken by parasites. This puzzle eventually played a central role in origin of life studies [11, 12].

In the eighties, a theoretical solution emerged, the Stochastic corrector model [13] inspired by ideas of group selection [14]. The idea is to use compartmentalization and selection to maintain functional replicators (ribozymes). The compartments in that model undergo cell division, which is a sophisticated feature that strongly constrains the allowed prebiotic scenarios.

In order to address this point, and also to assess the role of transient compartmentalization using a quantitative theoretical model, we introduced a general class of multilevel selection with transient compartmentalization [15]. This class includes several scenarios for the origin of life based on various types of compartments (lipid vesicles [16], pores [17, 18], inorganic compartments [19],‥) or various protocols of transient compartmentalization [20, 21] and a recent experiment, in which small droplets containing RNA in a microfluidic device [22] were used as compartments.

The related issue of cooperation between producers and non-producers has been discussed before [23]. When compartments are not well defined, their role can be played by spatial clustering, which can favor the survival of cooperating replicators [24, 25]. These ideas were combined in a recent study of a population of individuals growing in a large number of compartmentalized habitats, called demes [26]. Another recent related study on transient compartmentalization quantifies co-encapsulation effects in the context of directed evolution experiments [27].

In this paper, we go beyond the analysis carried out in our previous work [15] by including the effect of mutations and noise. The motivation of including mutations comes from experiments, since mutations play a role in the RNA droplet experiment [22] which inspired us. The motivation of discussing noise in more details comes from the realization that replication is inherently stochastic when a small number of replicators are present in compartments. Therefore, the deterministic approach used in our previous work [15] may seem like a severe approximation. In fact, it is important to appreciate that although our deterministic model neglects some sources of noise such as fluctuations in the growth rates, it includes fluctuations due to the smallness of the number of replicators present in the initial condition. This is a major source of noise because at the end of the exponential phase, the compartments no longer contain small numbers of molecules and therefore fluctuations become again negligible. A similar effect occurs when considering protein aggregation kinetics in small volumes: the fluctuations in the mass concentration of proteins at some later time are mainly due to the smallness of the number of proteins present in the initial condition [28].

In Sec. 2, we recapitulate the essence of the mutation-free model which we have introduced in [15], then in Sec. 3, we introduce an extension of that model to include deterministic mutations. The effect of noise is then covered in Sec. 4, which contains, in particular, in Sec. 4.1 a simple model for the replication of a single template by an enzyme, and in Sec. 4.4 an analysis of the growth noise in a population of such replicating molecules. The latter model is finally used to analyze the effect of noise on the transient compartmentalization dynamics introduced in the first section.

## 2. The mutation-free model

### 2.1 Definition of the model

We start from a large pool of molecules, which contains functional molecules called ribozymes and non-functional ones called parasites. Let the fraction of ribozymes in this pool be *x*. These molecules then seed a large number of compartments, which is treated in the infinite limit. A given compartment will contain *n* molecules, out of which *m* will be ribozymes and the remaining ones parasites. It follows that *n* is a random variable drawn from a Poisson distribution of parameter *λ*, while the number *m* follows a binomial distribution *B*_*m*_(*n, x*). The resulting probability distribution for seeded compartments is then

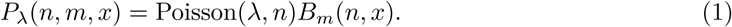

In addition to the replicating molecules, a large amount of replication enzymes *n*_*Qβ*_ and activated nucleotides *n*_*u*_ is supplied. We assume that all compartments contain the same amount of replication enzymes and activated nucleotides.

After seeding, the numbers of ribozymes *m* and parasites *y* grow exponentially, what in a deterministic model leads to

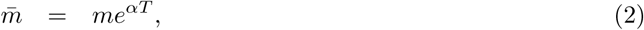

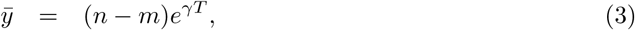

with *T* the time at the end of exponential growth phase, 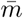 the number of ribozymes and 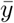 the number of parasites at time *T*. At the end of this growth phase, we have 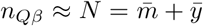, at which point further growth will be limited by the number of replication enzymes. As a result, after time *T*, the growth will be linear instead of exponential, but in any case, the system composition defined here by the relative fraction of ribozymes, will not change. For this reason, we focus on the final composition at time *T* which is controlled by the ratio Λ = *e*^(*γ-α*)*T*^. Here, we do not describe precisely the crossover between the exponential and the linear regime, which could be done using the notion of carrying capacity [29]. In that case, the growth would be described by logistic equations and the carrying capacity would be comparable to *n*_*Qβ*_. Note also that the exact time *T* may depend on *m, n*, but since in practice *n*_*Qβ*_ *⪼ m, n* this dependence has a small effect on the results of the model as we have checked in the Suppl. Mat. of Ref. [15].

In any case, the ribozyme fraction at the end of growth phase can be well approximated as

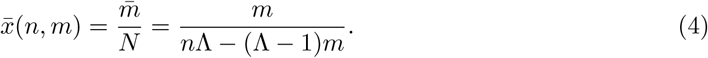

If parasites grow faster, we have *γ > α*, and thus Λ *>* 1, which is the regime considered in Ref. [15]. In Sec 3, we also consider regimes in which *γ < α*.

We now implement selection at the compartment level. In practice, selection could be autonomous or non-autonomous. For instance, in the experiment of Ref [22], the selection was non-autonomous: a measurement of the synthesis of a dye molecule by photodetection was used to promote or reject compartments according to the outcome of that measurement. Selection can in general be described by a selection function 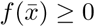. In our work, we have assumed that the selection function only depends on the final composition 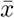 of the compartment.

For the ribozyme-parasite scenario, a natural choice for *f* is a monotonically increasing function of 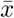. As an example, we will use the sigmoidal function

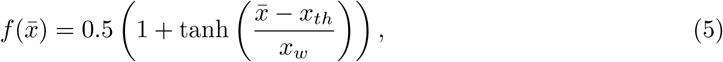

where *x*_*th*_ and *x*_*w*_ are dimensionless parameters, which describe respectively a threshold in the composition and the steepness of the function.

The compartments which have passed the selection step are then pooled together, forming a new pool of molecules from which future compartments can be seeded. The ribozyme fraction *x′* of this new ensemble is the average of 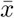 among the selected compartments

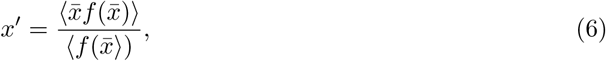

which is equivalent to

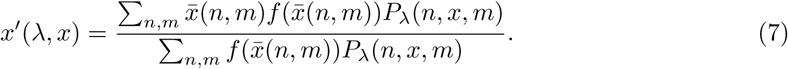

The transient compartmentalization cycle is then repeated, starting with the seeding of new compartments from that pool of composition *x′*.

Upon repetition of this protocol, the pool composition typically converges to a fixed point *x*^*\**^, which is a solution of

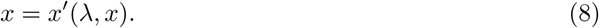

The stability of the fixed p oint *x* ^*\**^ changes when

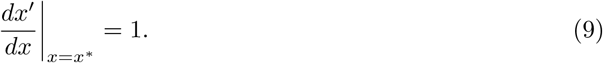

It is implicitly assumed that *x′*(*x*) is a sufficiently smooth function of *x* for this derivative to be defined.

### 2.2 Main dynamical regimes

Although finding a fi xed po int *x*^*\**^ is ge nerally di fficult, our ri bozyme-parasite mo del contains two simple fixed points: *x* = 0 a nd *x* = 1. By evaluating the s tability of these two fi xed po ints, four regimes can be distinguished, which are shown in the phase diagram in Fig 1. If *x* = 1 is stable and *x* = 0 unstable, ribozymes are stabilized and parasites are purged. If *x* = 0 is stable and *x* = 1 unstable, parasites deterministically invade the pool and purge ribozymes. If both *x* = 0 and *x* = 1 are unstable, trajectories from either side are attracted to a stable third fixed point 0 *< x* ^*\**^ *<* 1, leading to stable coexistence between parasites and ribozymes. Finally, if *x* = 0 and *x* = 1 are stable, their basins of attraction are separated by a third fixed point 0 *< x* ^*\**^ *<* 1, w hich i s unstable, and in this case we have a bistable regime in which the initial composition determines the fate of the system.

**Figure 1:**
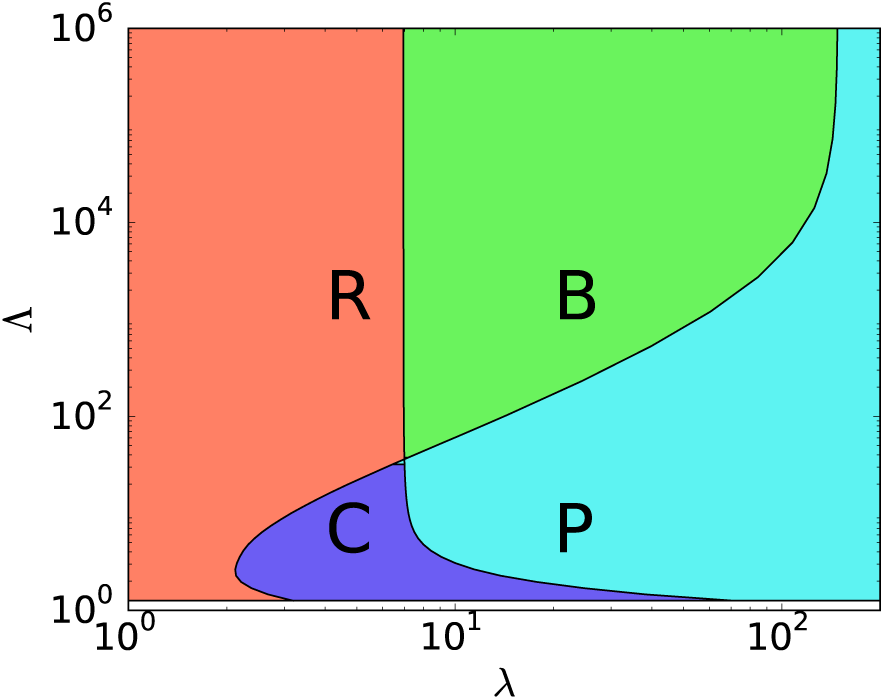
Original phase diagram of the mutation free model, taken from Ref[15]. The various phases are pure ribozyme (R), bistable (B), coexistence (C), pure parasite (P).

These conclusions can only be drawn provided there are not other fixed p oints b esides (*x* = 0*, x* = 1*, x* = *x*^*\**^). Extra fixed points come in pairs (one s table, o ne u nstable) a nd matter o nly if they are situated within (0, 1), in which case a stable coexistence and a bistable phase would be added to the behavior inferred from the other fixed p oints. For s imple monotonically increasing selection functions, we find t hat e xtra fi xed po ints are a ra re oc currence. Nevertheless a case where this occurs has been discussed in the Suppl. Mat. of Ref. [15].

### 2.3 Comparison to experiments

In addition to predicting the phase diagram associated with the long-time compositions reached by this transient compartmentalization dynamics, our theoretical model makes also predictions regarding the evolution of the ribozyme fraction as function of the round number, *i.e.* the number of completed cycles of compartmentalization. The model correctly reproduces that this fraction quickly goes to zero as function of the round number in bulk, less quickly with compartmentalization and no selection and even less quickly in the case of compartmentalization with selection. In the latter case, a finite fraction can be maintained for an infinite number of rounds provided *λ* is sufficiently small, corresponding to the coexistence region of the phase diagram.

In order to compare precisely the predictions of the model to the experiments of Ref. [22], it is important to know the value of key parameters such as Λ. Table 1 reports the experimental parameters measured in Ref. [22] for the ribozyme and three different parasites. The nucleotide length, its doubling time (*T*_*d*_), its relative replication rate (*r*) from which we infer Λ in the final column. The doubling time *T*_*d*_ for the ribozyme is related to the growth rate *α* by *T*_*d*_ = ln(2)*/α*, and similarly the doubling times of the parasites is *T*_*d*_ = ln(2)*/γ*.

**Table 1:** Lengths and doubling times for the parasites and ribozyme observed in Ref. [22], together with their relative aggressivity mesured by their relative growth rate *r*, and the corresponding values of Λ.

| Type | Length (nt) 2 | $T_d(s)$ | Relative $r$ | $\Lambda$ |
| --- | --- | --- | --- | --- |
| Ribozyme | 362 | 25.0 | 1.00 | 1 |
| Parasite 1 | 245 | 20.7 | 1.21 | 13 |
| Parasite 2 | 223 | 17.1 | 1.46 | 107 |
| Parasite 3 | 129 | 14.6 | 1.71 | 473 |

In the experiment, a typical compartment contains *λ* RNA molecules that can be ribozymes or parasites, 2.6 10^6^ molecules of Q*β* replicase, and 1.0 10^10^ molecules of each NTP. Replication takes place by complexation of RNA with Q*β* replicase, which uses NTPs to make a complementary copy. This copy is then itself replicated to reproduce the original. There is a large amount of nucleotides, so that exponential growth of the target RNA proceeds until *N ≈ n*_*Qβ*_. This large quantity of enzymes also means that in practice, the noise due to fluctuations in the number of enzymes should be very small. Starting from a single molecule, it takes *n*_*D*_ = log_2_ *n*_*Qβ*_ = 21.4 doubling times to reach this regime. In a parasite-ribozyme mixture, we can estimate Λ using the relative *r*:

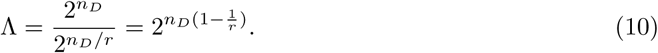

### 3. A modified model with deterministic mutations

In the deterministic model, we assume that a fraction *µ* of replicated ribozyme strands mutate into parasites. Thus, the equations describing the evolution of *m* and *y* in the growth phase assumes the form

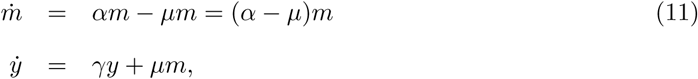

which yields for the first equation

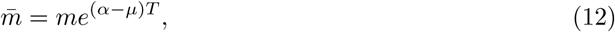

where 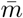 is again the number of ribozymes at the end of the growth phase and *m* the value at the initial time. Now substituting Eq. (12) into the equation for *y*, one finds

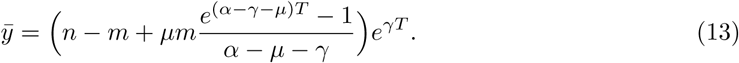

The ratio between the number of daughters of one parasite molecule and the number of daughters of a ribozyme molecule is now renormalized by the rate 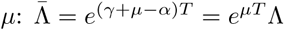, where Λ is the relative growth of parasites introduced previously in the mutation-free model.

The fraction of ribozymes at the end of the exponential phase is now given by

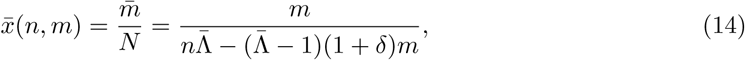

where *δ* = *µ/*(*α - µ - γ*). We call *δ* the mutation ratio, which is a dimensionless measure of mutation versus relative growth (competition). When *δ →* 0, we recover the mutation-free model, if *|δ| ⪼* 0 mutations become dominant.

Selected compartments are then pooled together, and the new average fraction of ribozymes becomes 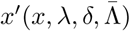. Note that for nonzero mutation rate (*µ >* 0), *x′* = 1 ceases to be a fixed point in this deterministic approach, since parasites will always appear at sufficiently long times. Therefore, the pure ribozyme (R) phase is no longer present in the phase diagram of fig. 2.

**Figure 2:**
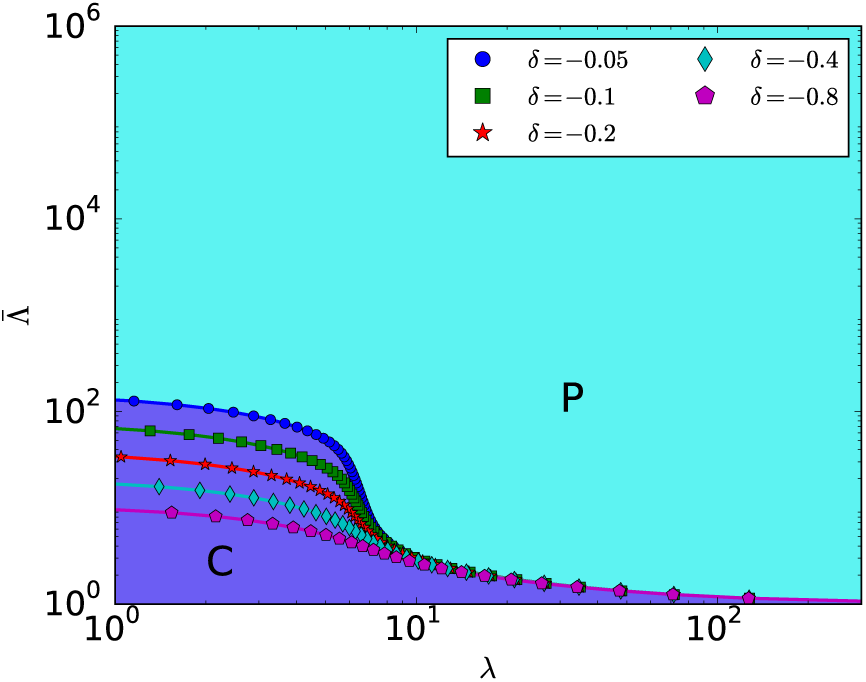
Phase diagram of the model with mutation in the case of prolific parasites. The selection function is given in Eq. (5). Phases are colored for *δ* = 0.05, other separatrices are plotted for various mutation strengths *δ*. The possible phases are coexistence (C), pure parasite (P).

The fixed point *x′* = 0 however is still present. If this fixed point is stable, we have a pure parasite phase. If it is unstable, there is stable coexistence at a fixed composition. If more fixed points appear, multiple stable compositions are in principle be possible.

### 3.1 The prolific parasites regime 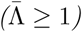

Prolific parasites have a better bulk reproductive success than ribozymes, when 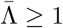, which is equivalent to *α ≤ µ* + *γ* and *δ <* 0. In a mutation-free model, this would imply necessarily a faster growth of parasites (*α < γ*), but in the present case, we could also allow for slower parasites as compared to ribozymes (i.e. *α > γ*), provided parasites are aided by a sufficiently high mutation rate *µ*.

The phase diagram is evaluated by testing the stability of the fixed point *x*′ = 0. We find an asymptote behaving like 1*/λ* for large *λ*, and plateaus for small *λ*. The ends of these plateaus locate in the limit *δ →* 0 at the position of the vertical line separating the ribozyme and bistable phase in the original phase diagram.

Let us first derive the right asymptote in the *λ* ≫ 1 limit. In this limit, we evaluate *x*′ by considering compartments of size *λ*

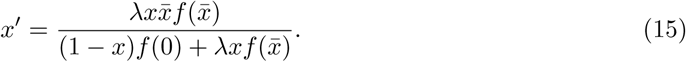

The fixed point stability condition *dx ′/dx′*|_*x*=0_ = 1 leads to

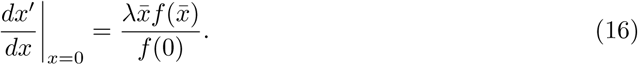

Upon substituting Eq. (14) evaluated at *m* = 1*, n* = *λ* and approximating 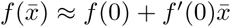, (for 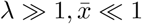) we find a quadratic equation for 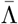, whose only physical solution 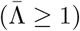 is

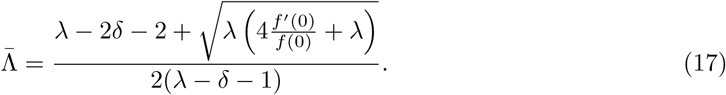

Since we consider monotonically increasing selection functions, *f′*(0) *>* 0. For *λ ≫ -δ*, we find

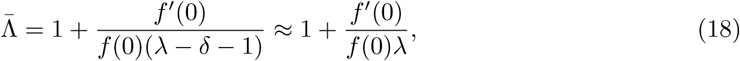

which is the same expression as the one found in the mutation-free phase diagram [15]. This explains why there is a single asymptote as *µ* is varied in the *λ ≫* 1 limit.

The plateaus extend to very low values of *λ*. We can find their location by considering only compartments of size *n* = 1. In that case, the final compositions can be 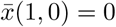 or

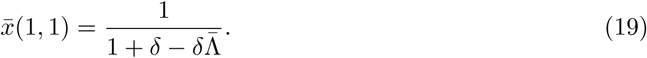

We then have for the composition recursion

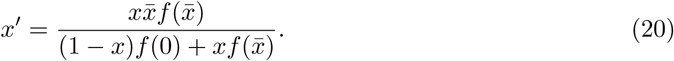

Evaluating the derivative of *x′*(*x*), we find

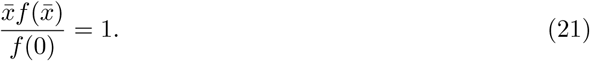

Substituting (19), we find that the location of plateaus obeys the implicit equation

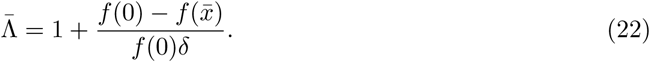

### 3.2 The prolific ribozymes regime 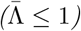

We now consider the opposite case where parasites are less prolific than ribozymes. This means *α ≥ µ* + *γ* and is equivalent to 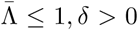. This implies that *α > γ* (less aggressive parasites) and is reminiscent of a quasipecies scenario in which a fit ribozyme succesfully outcompetes its parasites in bulk [10]. Since this can already happen in the absence of selection, we consider here the case where there is no selection, *i.e.*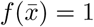

To analyze this regime we again assess the fixed point stability of *x′* = 0. We locate numerically the separatrix as shown in Fig 3. We obtain separatrices that for 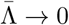 tend to a fixed value of *λ*.

**Figure 3:**
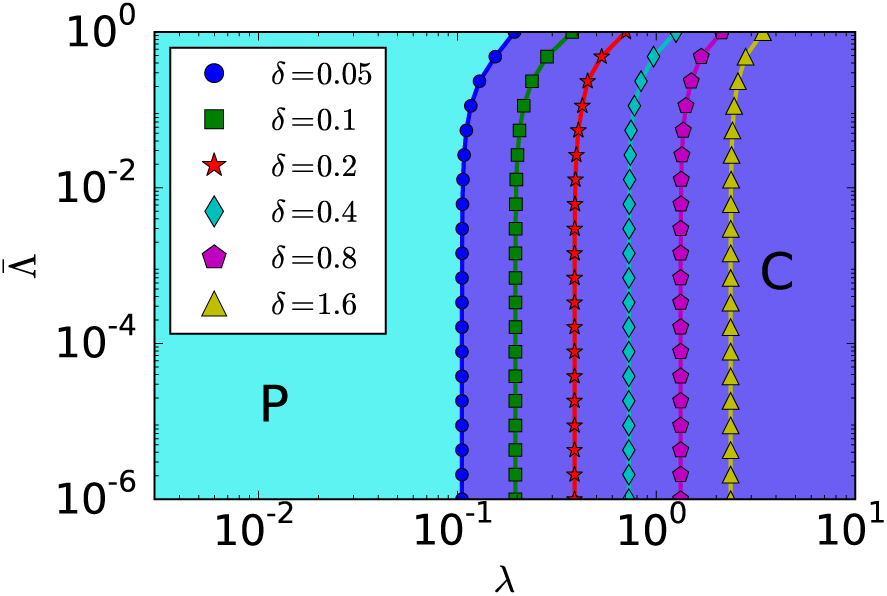
Phase diagram in absence of selection function for prolific ribozymes 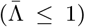. Phases are colored for *δ* = 0.05, separatrices are plotted for various mutation strengths *δ*. C: coexistence, P: pure parasite.

Let us start by observing that when 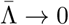, there are only two final compartment compositions for nonempty compartments: 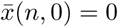 or 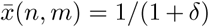 for *m >* 0. We can now distinguish between three initial compartment compositions: (i) only parasites, (ii) no parasites, no ribozymes, and (iii) containing at least one ribozyme. Their associated seeding probabilities are:

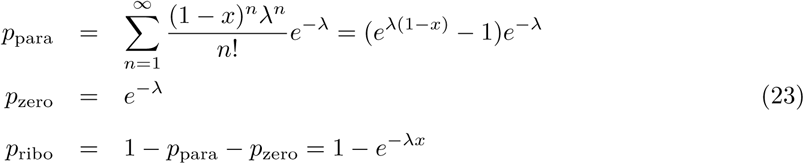

In that case, we can write the composition recursion equation as

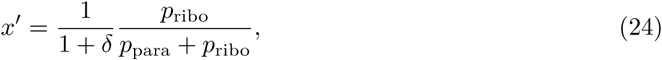

The condition *dx′/dx|*_*x*=0_ = 1 yields the expression

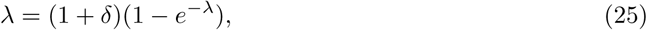

for the asymptote. For *λ ≪* 1, we obtain using (25)

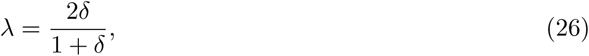

which agrees very well with Fig 3.

Notice that here the coexistence phase is located to the right of the asymptotes, and the parasite phase to the left, whereas in Fig 2 it is the other way around. An intuitive way to understand this is to consider the limit *λ →* 0. In this limit, nonempty compartments start with either a parasite or a ribozyme. The former will grow to a fully parasitic compartment, whereas the latter will contain ribozymes plus some parasites acquired by mutations. Therefore, at low *λ*, the ribozyme’s capacity to outgrow parasites (competition) cannot be exploited, leading to ribozyme extinction. It is only when ribozymes and parasites are seeded together that the differential growth rate becomes important, which becomes increasingly likely for higher *λ*. The phase boundaries in Fig. 3 mark the point where enough compartments engage in competition to allow for ribozyme survival. The mutation strength *δ* compares mutation rate to competition. When *δ →* 0, there is enough competition to ensure coexistence for all *λ*.

### 3.3 Error catastrophe

An error catastrophe corresponds to a situation where the accumulation of replication errors eventually causes the disappearance of ribozymes. Since there are only a parasite (P) and a coexistence phase (C) in the model with mutations, the error catastrophe means that the coexistence region shrinks at the benefit of the parasite phase as the mutation rate increases. One sees this effect in Fig. 2, which corresponds to the prolific parasites regime 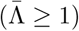 discussed above. In this figure, we see a larger coexistence region in the small *λ* region, because there the compartmentalization is efficient to purge parasites. As the mutation rate increases however, this region shrinks because the compartmentalization fails to purge the more numerous parasites.

In Fig. 4, a particular example is provided where *α* and *γ* are fixed, such that 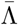 is fixed, and *µ* is varied. Since competition is fixed, we have *µ ∝ δ*. The resulting steady-state value *x* = *x*^*\**^ then decreases monotonically with *µ*, and reaches *x* = 0 when crossing the phase boundary in Fig 5. For small values of *λ*, this boundary corresponds to the plateau region, for larger values, this corresponds to the 1*/λ* asymptote. As can be seen in Fig 4, coexistence is stable for much higher values of the mutation rate *µ* when the compartment size *λ* is small. This means that compartmentalization with selection leads to a relaxed error treshold with respect to the bulk.

**Figure 4:**
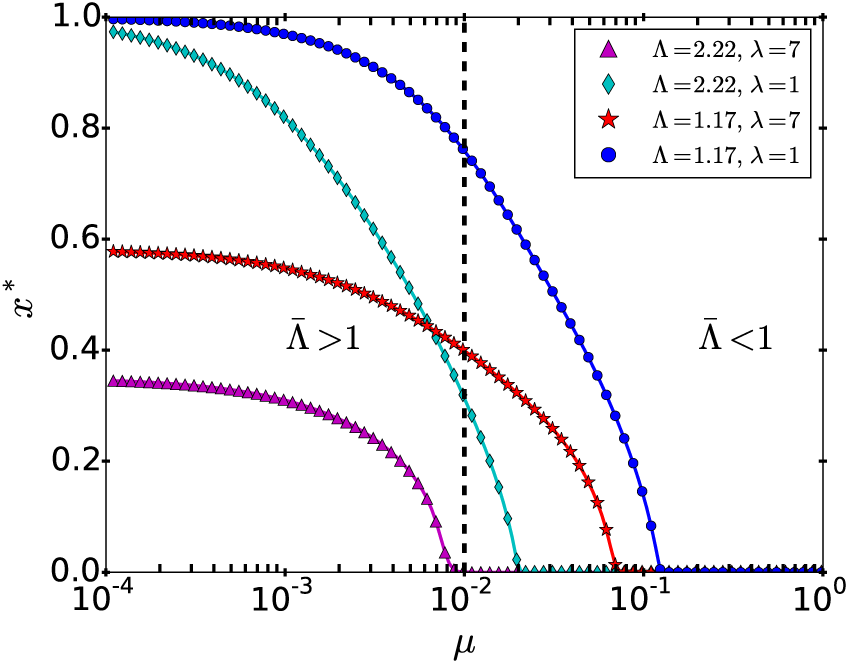
Steady state composition *x*^*\**^ as function of *µ*, *α* = 0.99*, γ* = 1.0. Critical rates *µ*^*\**^ corresponds to separation between P and C phases in Fig. 5.

**Figure 5:**
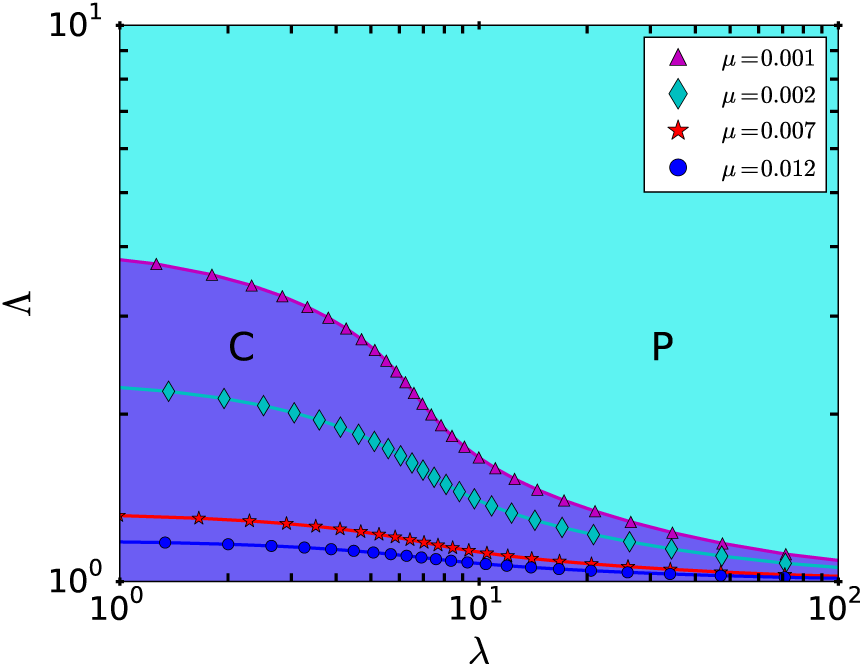
Phase diagram, drawn for f*α* = 0.99*, γ* = 1.0. Separatrices are drawn for *µ* values close to *µ*^*\**^ in Fig. 4, corresponding to an error catastrophe.

The error catastrophe was also studied in the absence of selection, and was shown to be in the prolific ribozymes regime 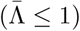. In Fig. 6, an example of this case is shown, and there too, we see that the steady-state value of the ribozyme fraction *x*^*\**^ decreases as *µ* is increased, until it reaches the phase boundary in Fig 7. In contrast to Fig. 4, where the error threshold decreases as the size of compartments increases, the trend is just the opposite in Fig. 6, which is expected since the role of ribozymes and parasites are exchanged here as compared to the prolific parasites regime.

**Figure 6:**
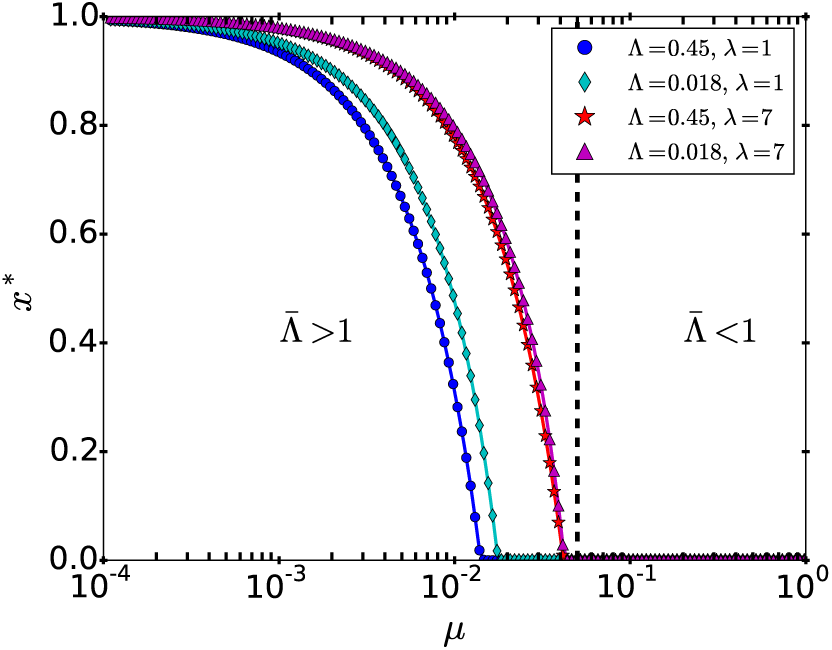
Steady state composition *x*^*\**^ as function *µ*, *α* = 1.0*, γ* = 0.95, in absence of selection (*f* = 1.0). Critical rates *µ*^*\**^ corresponds to separation between P and C phases in Fig. 7.

**Figure 7:**
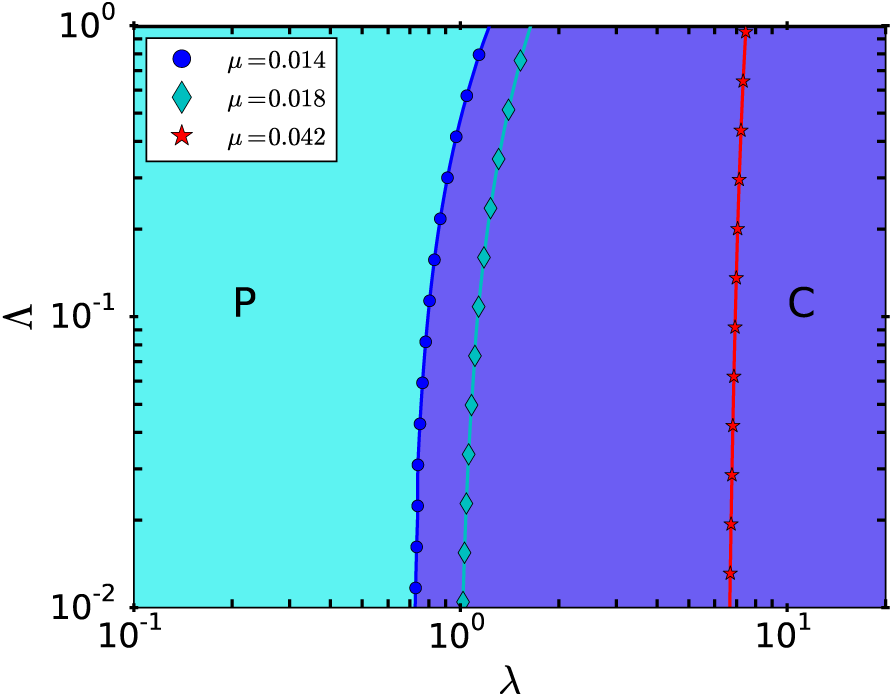
Phase diagram in absence of selection function (*f* = 1.0), drawn for *α* = 1.0*, γ* = 0.95. Separatrices are drawn for *µ* values close to *µ*^*\**^ in Fig. 6.

In the prolific parasites regime, 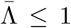 with selection, it is interesting to recast the error threshold as a constraint on the length of a polymer to be copied accurately, as done in the original formulation of the error threshold [10]. Let us introduce the error rate per nucleotide, ϵ. Then, for a sequence of length *L*, we have *α - µ* = *α*(1 *- ϵ*)^*L*^. Since ϵ *≪* 1, it follows from this that *µ* = *αϵL*. When *α ≃ γ*, we have ln 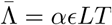. Using Eq. (22), we find that the condition to copy the polymer accurately is

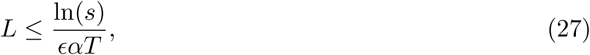

where 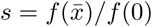 and *αT/* ln 2 is the number of generations. This criterium has a form similar to the original error threshold [10], namely

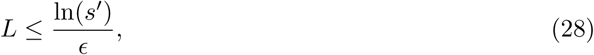

where *s′* = *α/γ* represents the selective superiority of the ribozyme. In our model, the equivalent of *s′* is *s* which characterizes the compartment selection.

## 4. Noise in growth

For deterministic growth, given by Eqs. (2)-(3), fluctuations in the growth rates, denoted *α* for the ribozymes and *γ* for the parasites, have been neglected. In order to estimate whether fluctuations in the growth rates of ribozymes and parasites could have a strong effect, we introduce in the next section a model for the replication process of such molecules by a replicating enzyme. This model includes noise due to the stochastic binding of the replicating enzyme to template molecules and the noise due to the stochasticity of monomer addition once the enzyme is bound to a template. Importantly, this model assumes that the replicase once bound stays always active until completion of the copy of the template, therefore the possibility that the replicase falls off the template before completion of the copy is neglected. Similarly, any effects associated with the interaction of multiple replicases on the same template are neglected. In fact, when the replicase falls off of its template, the copying process is aborted and the shorter chain which has been produced in this way becomes a parasite. We can therefore describe such a process as a mutation using the framework of the previous section. To separate the effects due to mutations and noise clearly, we disregard from now on the possibility of mutations, and we focus in the following on the description of the noise associated with replication. Such a noise can stabilize the ribozyme phase at the expense of coexistence, and the coexistence phase at the expense of the parasite phase. The noise of replication becomes very small when the rate-limiting step is nucleotide incorporation, in which case one can use a deterministic approach.

### 4.1 A minimal model for the replication process

The replication of an RNA strand *A* by a replicase can be considered to proceed through two stages. In the first stage, an RNA molecule *A* binds to a polymerase *E*, to form a complex *X*_0_

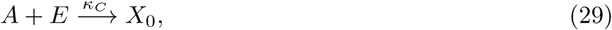

with the rate *κ*_*C*_.

Subsequently, activated nucleotides *X* are incorporated in a stepwise fashion to the complementary strand. A complex of *E* and *A* with a complementary strand of length *n* will be denoted by *X*_*n*_, and the strand grows until the final length *L* is achieved, such that

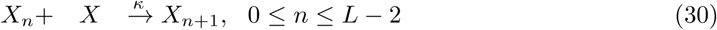

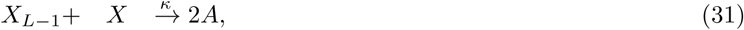

where for simplicity we have assumed the same rate *k* for both reactions. Let us denote by *t* the total time to yield 2*A* from *A*, which is the sum of the time associated with the step of complex formation, *t*_*C*_ and with the step of *L* nucleotide incorporations *t*_*L*_. We thus have

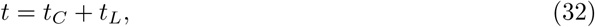

with 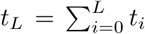 and *t*_*i*_ the time for adding one monomer, which we assumed is distributed according to

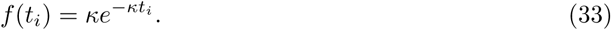

For simplicity, we choose a single value *κ* for all monomer additions. The time for the formation of the complex, *t*_*C*_ is similarly distributed according to

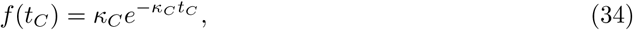

where *κ*_*C*_ = 1*/(t*_*C*_ ).

Let us denote the moment generating function of *t*_*C*_ by *M*_*C*_ (*s*) and similarly for *t*_*L*_ by *M*_*L*_(*s*) with:

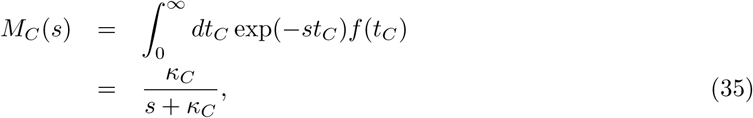

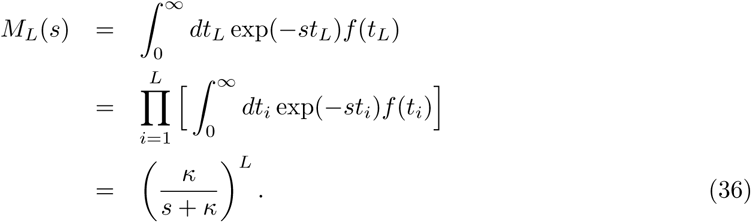

From *M*_*L*_ one obtains the distribution of replication time *f* (*t*_*L*_) by performing an inverse Laplace transform:

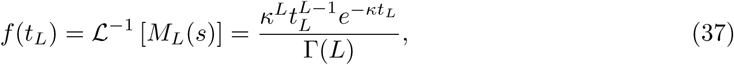

where ℒ ^*-*1^ represents the inverse Laplace transform. This equation shows that the replication time distribution of one strand of length *L* follows a Gamma distribution [30]. For *L* = 1, Eq. (37) becomes a simple exponential distribution, which is a memoryless distribution. For *L >* 1, this distribution has memory and the growth in the number of RNA strands can no longer be described as a simple Markov process. Note that the Gamma distribution is peaked around the mean value of *t*_*L*_, namely *L/κ* for *L ≫* 1. In this limit, the replication time has very small fluctuations. This feature has recently been exploited to construct a single-molecule clock, in which the dissociation of a molecular complex occurs after a well-controlled replication time[31].

### 4.2 Coefficient of variation of the replication time

Let us now study the coefficient of variation of the full time *t*, which includes the diffusion of the replicase and the replication step. The generating function of *t* is clearly *M* (*s*) = *M*_*D*_(*s*)*M*_*L*_(*s*). Thus, the cumulant-generating function defined as *K*(*s*) = ln *M* (*s*), yields the two moments of the distribution of *t*, namely the mean *(t)* and the variance 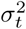. We have

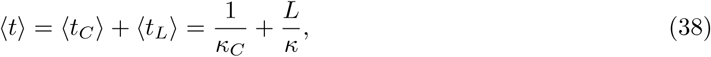

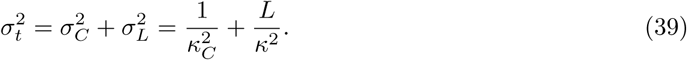

Thus the coefficient of variation of the replication time, namely *σ*_*t*_*/(t)* is given by

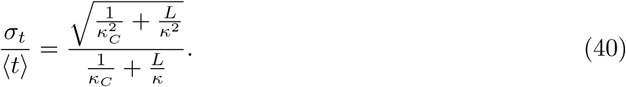

Fig 8 shows this quantity as function of the length *L* and of the ratio of the rates (*κ*_*C*_ */κ*).

**Figure 8:**
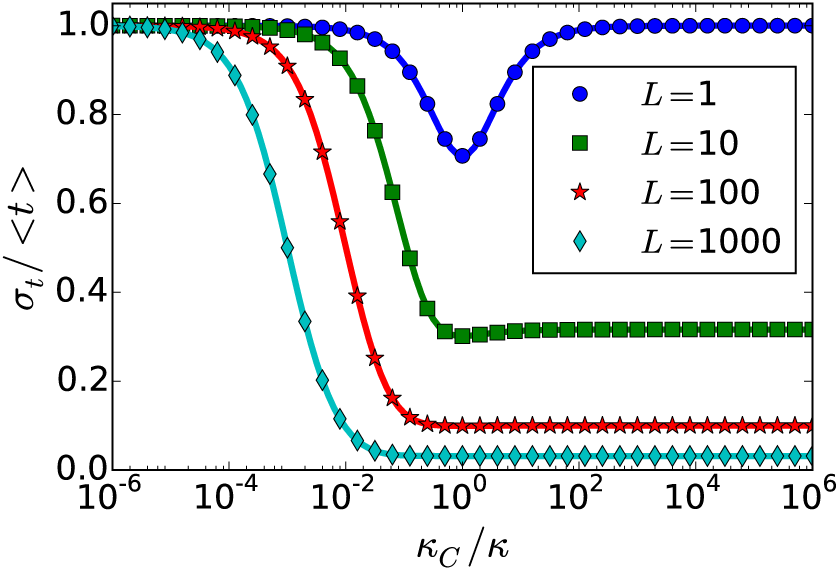
Waiting time variability *σ*_*t*_ */ 〈t〉* for various polymer lengths *L*, as a function of the ratio of typical times for replication and complex formation

There are two regimes: on one hand, when *L/κ ≫* 1*/κ*_*C*_, the time taken by the replication step dominates over the time for the replicase to diffuse to its target. If in addition 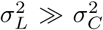, the coefficient of variation of the time *t* scales as 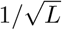 and therefore becomes very small for long strands. This power-law regime in indeed visible as plateaus in Fig 8.

On the other hand, when 1*/κ*_*C*_ *≫ L/κ*, the time to form a complex between the replicase and its template dominates over the replication time. This regime has a large coefficient of variation since *σ*_*t*_ *≃ 〈t〉* as also seen in Fig 8.

### 4.3 The generations representation

Let us look at these two growth regimes in a generations representations, where by generations we mean an event of copy of the template by the replicase. The diffusion-limited regime corresponds to Fig. 9, while the replication-limited regime corresponds to Fig. 10. In this representation, the differences in the two growth regimes become very clear. In the replication-limited regime, generations remain synchronized, until enough noise has accumulated over multiple generations. For two independent strains, generations become desynchronized after about 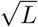 generations. In contrast, in the diffusion-limited regime, fluctuations are very large due to lack of synchronicity in the growth.

**Figure 9:**
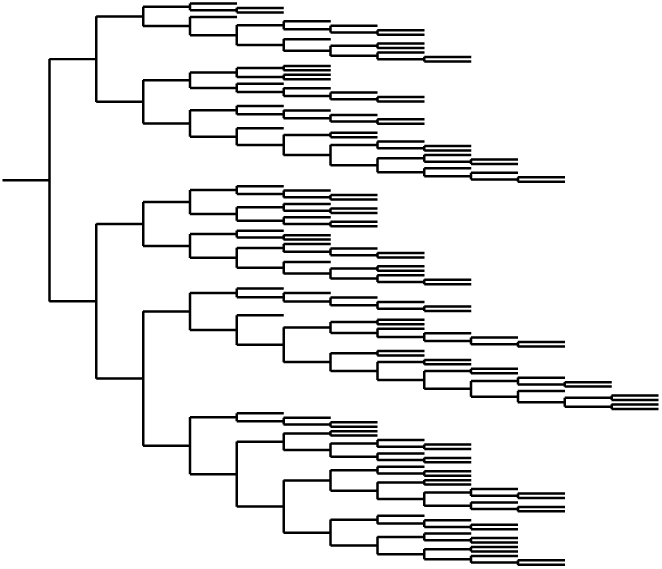
Generations representation of growth in the diffusion-limited regime. The simulation ends when the population size has reached 128. The horizontal axis corresponds to the generation number.

**Figure 10:**
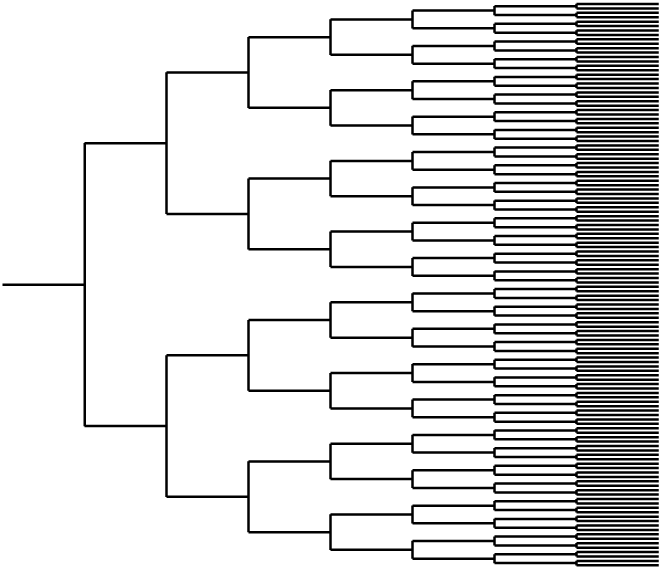
Generations representation of growth in the replication-limited regime. The simulation ends when the population size has reached 128. The horizontal axis corresponds to the generation number.

These figures have been obtained by simulating the growth of a replicating mixture starting from a single strand. The simulation follows *k* RNA-enzyme complexes, and for each the variable *n*_*k*_ measures the length of the growing complementary strand. For every nucleotide incorporation event, a strand *i* is chosen with probability 1*/k*, after which its number of nucleotides is updated from *n*_*i*_ to *n*_*i*_ + 1. When *n*_*i*_ + 1 = *L*, we set *n*_*i*_ = 0, we update *k* to *k* + 1, and then we introduce an extra strand variable *n*_*k*+1_ for the new strand. Both the replication-limited regime and the diffusion-limited regime can be modeled using this simulation. In the latter case, we need to choose *L* = 1, which corresponds to exponentially distributed replication times.

### 4.4 Population-level noise

In sec 4.2, we have analyzed the noise associated with the replication of a single strand. Ultimately, we wish to quantify the compositional variation of the final population. In order to do so, we turn to the theory of branching processes with variable lifetimes taken randomly from a fixed distribution [32]. As explained in AppendixA, this framework describes theoretically a population that grows exponentially starting from a single individual. In our molecular system, this single individual plays the role of the single molecule present in the initial condition before the replication starts; while the distribution of the lifetimes is the replication time distribution *f* (*t*_*L*_) obtained in Eq (37).

For *t*_*L*_ *≫ L/κ*, we find that the average population (starting from a single individual) *µ*^(1)^ scales as 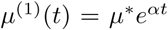, with a growth rate *α ≃ κ* ln(2)*/L*. The coefficient of variation of the population size *σ*^(1)^*/µ*^(1)^ is

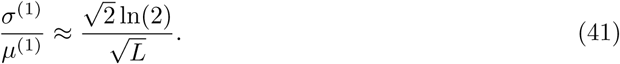

The renewal theory on which these results are based, can be generalized to the case that there are *n* individuals in the initial condition as shown in AppendixB. The full solution is rather complicated due to correlations between the subpopulations generated by the different molecules present in the initial condition. In the following, we neglect these correlations: therefore the *n* initial molecules generate *n* independent subpopulations, which all start at size 1 and follow the branching process described above and in AppendixA. In that case, each subpopulation now has a mean *µ*^(1)^ = *µ*^(*n*)^*/n* and a standard deviation 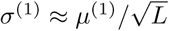. This then allows to write

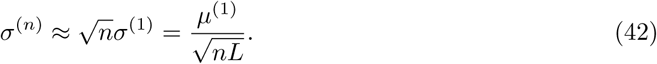

We show in Fig. 11 that the corresponding coefficient of variation, *σ*^(*n*)^*/µ*^(*n*)^, agrees well with simulations of the branching process. The 2000 simulation runs were stopped after a time *t*^*\**^ such that 〈*N* (*t*^*\**^)〉 *≃* 5000.

**Figure 11:**
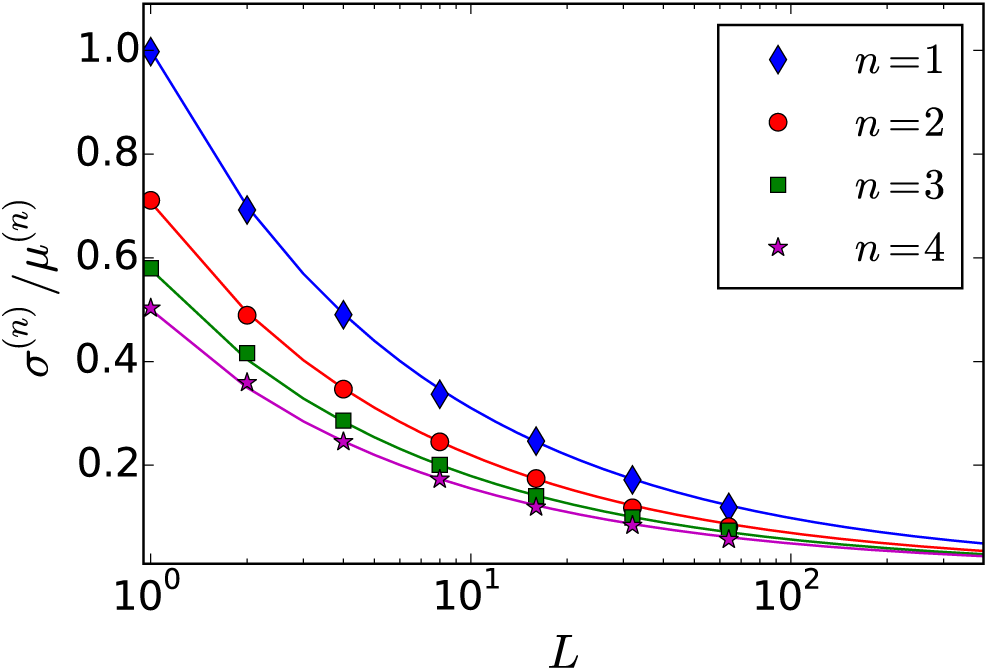
Coefficient of variation of the population size *N* as function of the initial population size *n*. The <u>re</u>sults have been averaged over 2000 runs. The solid lines represent the theoretical prediction: 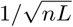.

### 4.5 Fluctuations in logistic growth

The problem of two species competing for the same resources has been studied in the literature and offers a complementary perspective on the role of noise in a growing population, which has been studied in the previous section. Let us consider two such species, which typically start with a few individuals and then grow according to logistic noise. As shown in Ref. [29], when the carrying capacity is reached, the number of each species is subject to giant fluctuations (the coefficient of variation is of the order of unity) when the two species have similar growth rates. In the terminology introduced in previous section, this model applies to the diffusion-limited regime (*L →* 1), where a Markov description of the population dynamics is applicable.

Keeping the notations of the first section, we denote by *n* the initial number of molecules, which splits into *m* ribozymes and *y* parasites, and by *N* the final number of molecules in the compartment. In the neutral case (*α* = *γ*), the moments of the number of ribozymes 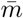 are found to be [29]:

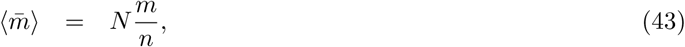

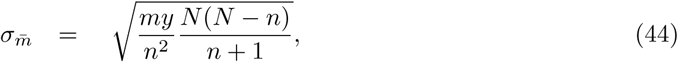

with again *y* = *n - m*. Since *N* remains fixed,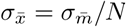. This means that

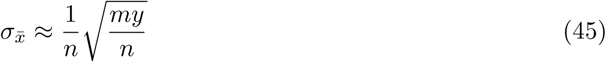

for *N ≫n*, which means that the noise in the composition depends primarily on the number of individuals in the initial condition. Let us denote *s* = *α/γ -* 1 ≪ 1, with *s ≪* 1 and *ρ* = ln(*N/n*). In Ref. [29], it was shown that

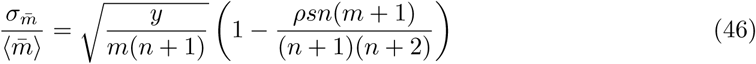

In general, the dynamics of the composition has a large variability for: (i) small compartments (*n ∼ O*(1)), (ii) mixed compartments (*m, y >* 0), and for *m ≈ y*, (iii) comparable growth rates (*s →* 0).

Such a coefficient of variation is asymptotically constant on long times and the constant only depends on the initial number of molecules. A similar scaling for the coefficient of variation holds in a number of other physical situations, such as for the fluctuations in the number of protein filament formed in small volumes [28].

### 4.6 Noise in transient compartmentalization

Let us now apply the results of the section 4.4 to analyze the effect of the growth noise on our transient compartmentalization dynamics. Let us assume that the length of the ribozymes is *L*_*α*_ and that of the parasites *L*_*γ*_. For experimental values of these parameters we refer the reader to Table 1. In section 2, we have defined *m, y* to be the initial number of ribozymes and parasites and 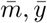 to be the final mean number of ribozymes and parasites at the end of the growth phase in a given compartment. Using Eqs. (41)-(42), we obtain

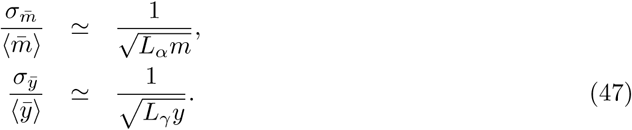

Since the ribozyme fraction 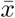 at the end of the exponential phase is given by 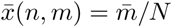 and *N ≃ n*_*Qβ*_, the noise on 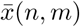 takes the following form:

**Figure 12:**
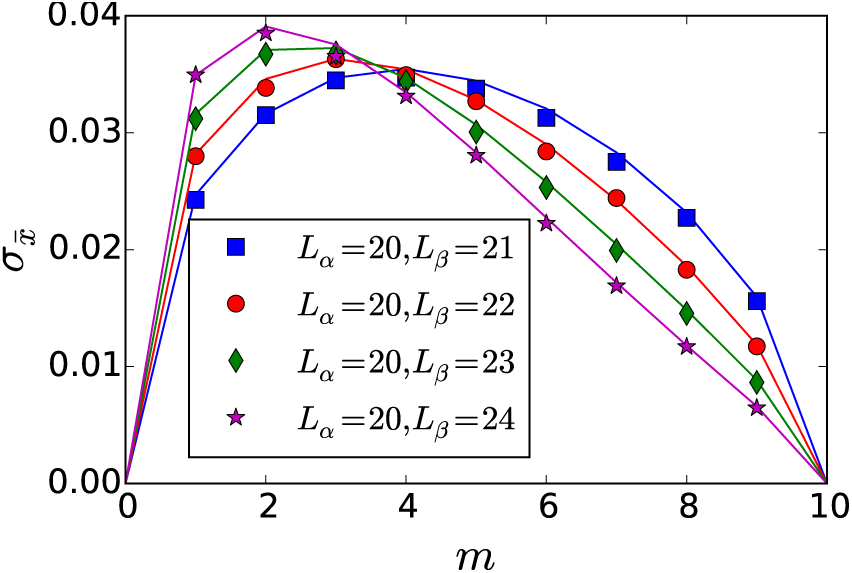
Standard deviation of the ribozyme fraction, 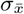, as predicted from simulations (symbols), and compared with predictions from Eq. (48) (solid lines). For each initial composition (*m, n*), 10000 simulations were performed until a time *t*^*\**^ such that 〈*N* (*t*^*\**^)〉 *≃* 5000 and by choosing *α/γ* = *Lγ /Lα*.

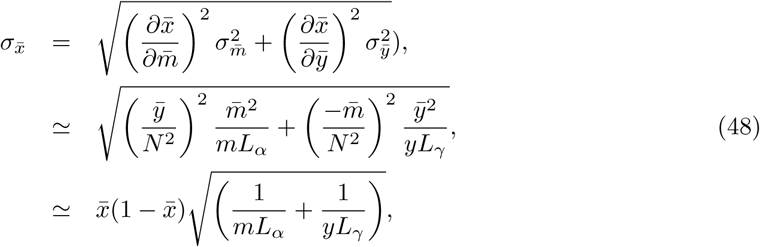

where we have used Eq. (42) with 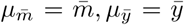. The factor 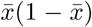 is largest for 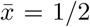 and vanishes for pure parasite and pure ribozyme compartments, which means that this noise can be neglected when Λ ≫ 1 or Λ ≪ 1. Note that if we choose *α* = *γ* (and thus 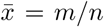), and *L*_*α*_ = *L*_*γ*_ = 1, Eq. (48) becomes

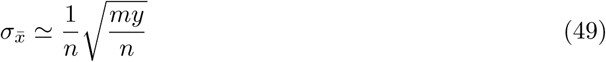

which is consistent with Eq. (45) which was found using a different formalism[29].

Using the parameters of Table 1 and (41), we can quantify the level of noise in the number of ribozymes or parasites. We find from this table that the ribozyme size was *L* = 362, and that the experiment should be in the replication-limited regime because the diffusion time scale should be approximately over 2 *·* 10^4^ times smaller than replication times of the order of 10s. The noise in composition should be maximal when we start with one ribozyme and one parasite of equal length, and with *α* = *γ*, which on average gives 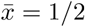. Consequently, the noise in composition is at most 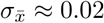.

### 4.7 A weak noise approach

The growth equations given by Eqs. (2) and (3) are deterministic in nature, which means that a given initial condition (*n, m*) yields a unique final composition 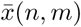. In contrast to that in a stochastic approach, a given *n* and *m* lead to many different trajectories, which means that 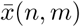 is a random variable with a probability distribution 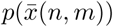. Consequently, the ribozyme fraction after one round is

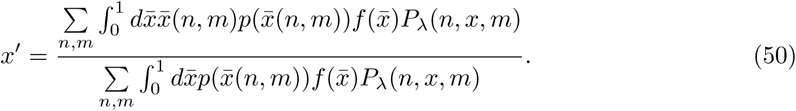

This expression is computationally demanding to evaluate for *λ ≫* 1, but it can be simplified significantly in the weak noise limit.

In order to construct a phase diagram in this limit, we simplify Eq. (50), by considering 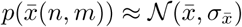, where 𝒩 denotes a normal distribution with mean 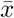 and standard deviation defined by Eq. (48). From Eq. (48) we expect the effect of noise to be largest when *λ, L* and Λ are close to 1 (if 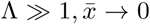). In Fig. 13, the original phase diagram from Ref. [15] is shown together with the modified phase boundaries (dotted lines) due to the presence of Gaussian noise using Eq. (50) for the case that *L*_*α*_ = *L*_*γ*_ = 3.

**Figure 13:**
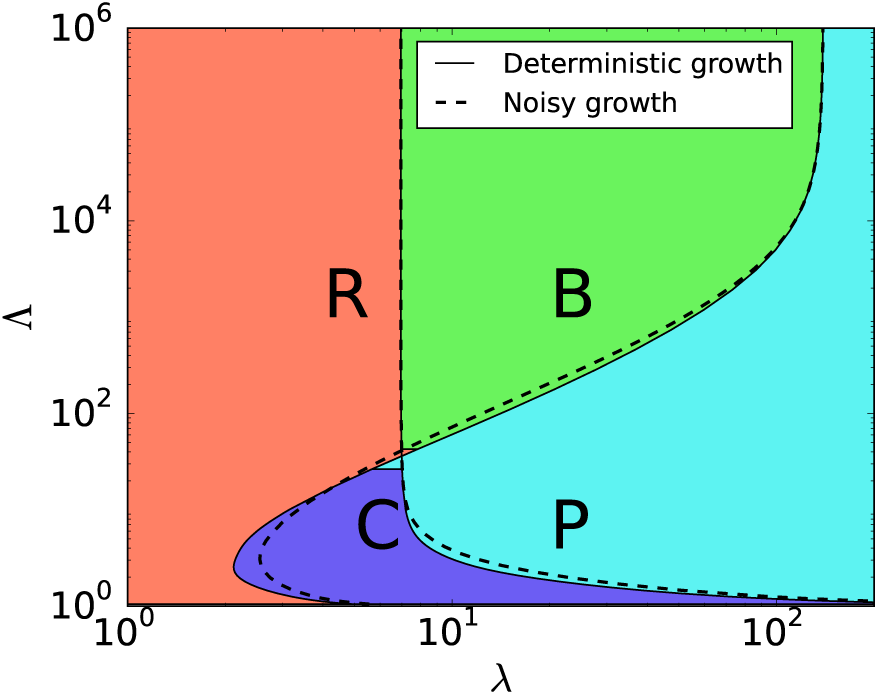
Phase diagram for ribozyme-parasite scenario in presence of noise given by Eq. (48), for *Lα* = *Lγ* = 3.

Given that the amplitude of this type of noise should rapidly diminish for larger *L*, and that *L ∼ O*(100) in the experiment, we expect our ribozyme-parasite scenario to be well-described by a deterministic dynamics. We also see that the noise stabilizes the pure ribozyme phase (R) with respect to the coexistence phase (C) because in the presence of noise, the R region has grown at the expense of the C region. Similarly, the noise stabilizes the coexistence region (C) against the parasite region (P).

## 5. Conclusion

In this paper, we have carried out two important extensions of our previous work on transient compartmentalization [15], by including the effect of mutations and noise in such systems. This new study confirms one result of our previous work, namely that transient compartmentalization alone can stabilize functional replicators in the absence of a division of compartments of the kind considered in the Stochastic corrector model [13]. We can now add to that, that this property is robust with respect to mutations and noise, an important aspect for Origin of life studies.

In the presence of mutations, we have found that the phase diagram of long-time composition of this system only contains the parasite and the coexistence phases. The case where ribozymes grow faster than the parasites can be analyzed in terms of a modified error threshold, which interestingly now depends on the dynamics of compartmentalization and selection.

In order to analyze the role of noise in this system, we have introduced a simple model for the replication of a template by an enzyme. In the replication limited regime of that model, which should correspond to our experimental conditions, a low noise should be present at the population level, which we have quantified using tools borrowed from the theory of branching processes. In the end, we have studied the modified phase diagram of our model in this weak noise limit.

Of course, the two effects that we have studied here separately, namely mutations and noise, could be present simultaneously. We cannot also exclude that a more detailed modeling of the molecular replication or a different form of compartmentalization dynamics could lead to features not captured by the present treatment. Nevertheless, we think that the present framework represents a basis on which further studies could be built. We hope that our work may not only contribute to studies on the Origin of life but also to future developments on related important experimental techniques such as digital quantitative PCR [33] or Directed Evolution [34].

In order to analyze the role of noise in this system, we have introduced a simple model for the replication of a template by an enzyme. In the replication limited regime of that model, which should correspond to our experimental conditions, a low noise should be present at the population level, which we have quantified using tools borrowed from the theory of branching processes. In the end, we have studied the modified phase diagram of our model in this weak noise limit.

Of course, the two effects that we have studied here separately, namely mutations and noise, could be present simultaneously. We cannot also exclude that a more detailed modeling of the molecular replication or a different form of compartmentalization dynamics could lead to features not captured by the present treatment. Nevertheless, we think that the present framework represents a basis on which further studies could be built. We hope that our work may not only contribute to studies on the Origin of life but also to future developments on related important experimental techniques such as digital quantitative PCR [33] or Directed Evolution [34].

## Acknowledgements

A.B. was supported by the Agence Nationale de la Recherche (ANR-10-IDEX-0001-02, IRIS OCAV). L.P. acknowledges support from a chair of the Labex CelTisPhysBio (ANR-10-LBX-0038). We would like to thank Y. Rondelez for many important and insightful suggestions. We acknowledge stimulating discussions with B. Houchmandzadeh.

## AppendixA Population-level noise generated by a single individual in the initial condition

Let us consider an age-dependent renewal process, in which the probability density of branching at age *t* is given by *f* (*t*), and upon branching, the probability of having *k* offspring is given by *ϕ*_*k*_ (assumed to be age-independent for simplicity). We would like to evaluate the behavior of the number *N* (*t*) of individuals at time *t*. Let us define the function *h*(*s*) by

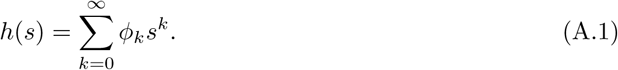

We also define the generating function for the process *N* (*t*) by

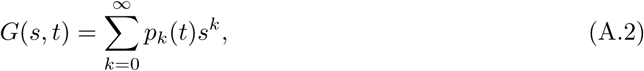

where *p*_*k*_(*t*) is the probability that *N* (*t*) = *k*. We assume that 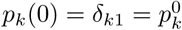, i.e., that we start from a single object. We can then evaluate *p*_*k*_(*t*) by adding the probability that no branching has occurred between 0 and *t*, which is given by 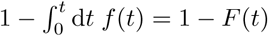, with the effect of the first branching at time ^0^*u*,such that 0 *< u < t*. We obtain

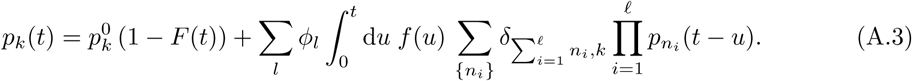

Multiplying by *s*^*k*^ and summing, we obtain

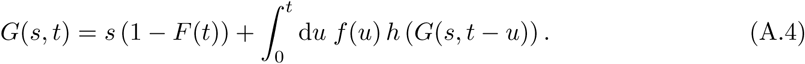

Taking the derivative with respect to *s* at *s* = 1, we obtain the following equation for the average *µ*(*t*) = Σ_*k*_ *k p*_*k*_(*t*):

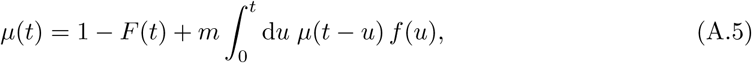

where *m* = *h ′* (1) =Σ _*k*_ *k ϕ*_*k*_ is the average number of daughters upon branching.

To solve this equation in the limit *t → ∞*, let us multiply both sides by e^*-αt*^ and take the limit. Since lim_*t→∞*_ *F* (*t*) = 1, we obtain

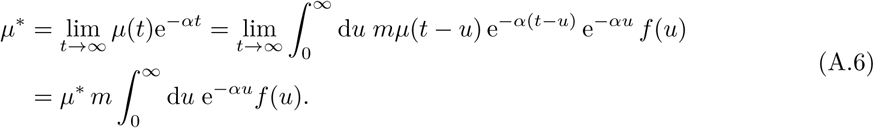

This equation allows for a solution different from 0 and *∞* if *α* is chosen to satisfy

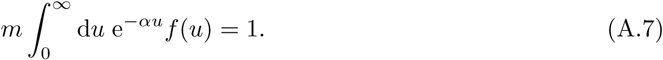

Then, making use of a result by Smith [**?**], we obtain

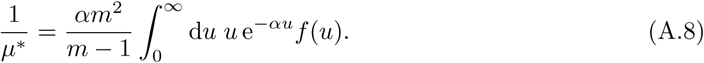

As a consequence, we have

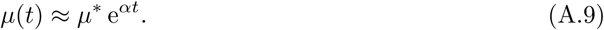

In the case we are considering we have

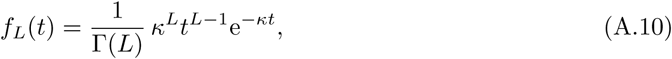

and *m* = 2, which yields

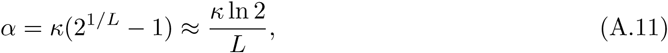

giving, as long as *L »* 1,

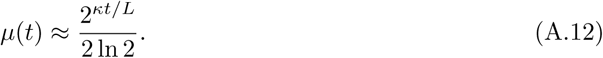

We can use this framework to also evaluate higher moments of the population size, and from that obtain the coefficient of variation of the population size which characterizes the amplitude of the noise. Let us denote the second derivative of the generating function with respect to *s* by *ζ*

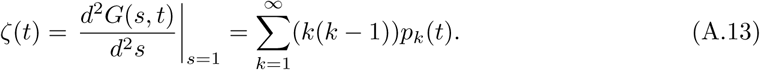

At large times, *ζ*(*t*) *≈ ζ* e*^2*αt*^. The variance of the population size *σ*^2^ follows from the standard relation:

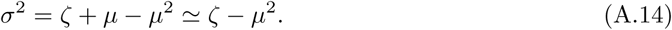

For the specific case we are considering, we find

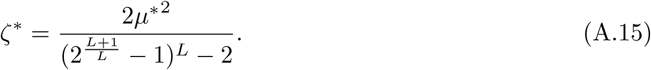

After extracting the leading contribution in the large *L* limit, we find:

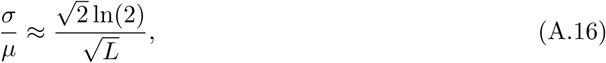

which is numerically close to 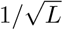 since 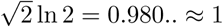.

## AppendixB Population-level noise generated from *n* individuals in the initial condition

If we start from *n* individuals rather than just one, we can write the probability to have *k* individuals at time 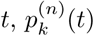., in terms of the subpopulations generated by *n* single individuals,

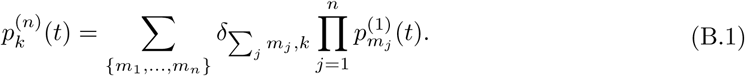

Here, 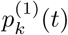 denotes the probability of having a population size of *k* at time *t*, starting from one individual, which was considered in AppendixA. Note that we have added an addition superscript (1) to the notation used in AppendixA to emphasize the initial condition. From this equation, the new generating function follows:

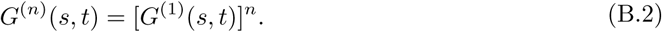

From this equation, we obtain the average,

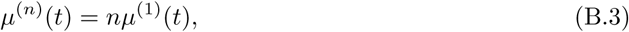

which expresses the average with *n* initial strands in terms of the average with one initial strand. For the second moment, we obtain

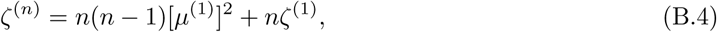

We can then extract *σ*^(*n*)^ by using Eq. (A.14), which yields

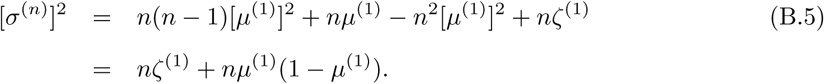

Together with Eq. (B.3), this leads to

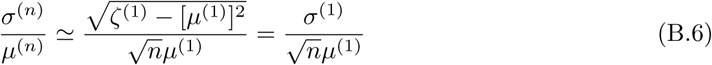

which is the coefficient of variation found previously for a single individual in the initial condition, divided by 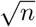 as expected for the growth from independent individuals. This confirms the scaling found in Eq. (42).

## References

[1] A. Jaiman, M. Thattai, Algorithmic biosynthesis of eukaryotic glycans, BioRxiv doi:10.1101/ 440792.

[2] A. I. Oparin, Origin of Life, Dover, 1952.

[3] C. P. Brangwynne, C. R. Eckmann, D. S. Courson, A. Rybarska, C. Hoege, J. Gharakhani, F. Jülicher, A. A. Hyman, Germline p granules are liquid droplets that localize by controlled dissolution/condensation, Science 324 (5935) (2009) 1729–1732. doi:10.1126/science.1172046.

[4] D. Zwicker, R. Seyboldt, C. A. Weber, A. A. Hyman, F. Jülicher, Growth and division of active droplets provides a model for protocells, Nat. Phys. 13 (4) (2017) 408–413. doi:10.1038/nphys3984.

[5] E. Brinke, J. Groen, A. Herrmann, H. A. Heus, G. Rivas, E. Spruijt, W. T. S. Huck, Dissipative adaptation in driven self-assembly leading to self-dividing fibrils, Nature Nanotechnology 13 (9) (2018) 849–855. doi:10.1038/s41565-018-0192-1.

[6] K. K. Nakashima, J. F. Baaij, E. Spruijt, Reversible generation of coacervate droplets in an enzymatic network, Soft Matter 14 (2018) 361–367. doi:10.1039/c7sm01897e.

[7] P. G. Higgs, N. Lehman, The RNA world: molecular cooperation at the origins of life, Nat. Rev. Genet. 16 (1) (2015) 7–17. doi:10.1038/nrg3841.

[8] F. Dyson, Origins of life, Cambridge University Press, 1985.

[9] S. Spiegelman, I. Haruna, I. B. Holland, G. Beaudreau, D. Mills, The synthesis of a self-propagating infectious nucleic acid with a purified enzyme, Proc. Natl. Acad. Sci. USA 54 (3) (1965) 919–927. doi:10.1073/pnas.54.3.919.

[10] M. Eigen, Self-organization of matter and the evolution of biological macromolecules, Naturwissenschaften 58 (10) (1971) 465–523. doi:10.1007/BF00623322.

[11] J. Maynard Smith, E. Szathmáry, The Major Transitions in Evolution, Freeman, Oxford, 1995.

[12] N. Takeuchi, P. Hogeweg, Evolutionary dynamics of rna-like replicator systems: A bioinformatic approach to the origin of life, Physics of Life Reviews 9 (3) (2012) 219–263. doi:10.1016/j. plrev.2012.06.001.

[13] E. Szathmáry, L. Demeter, Group selection of early replicators and the origin of life, J. Theor. Biol 128 (4) (1987) 463–86. doi:10.1016/S0022-5193(87)80191-1.

[14] D. S. Wilson, A theory of group selection, Proc. Natl. Acad. Sci. USA 72 (1) (1975) 143–146. doi:10.1073/pnas.72.1.143.

[15] A. Blokhuis, D. Lacoste, P. Nghe, L. Peliti, Selection dynamics in transient compartmentalization, Phys. Rev. Lett. 120 (2018) 158101. doi:10.1103/PhysRevLett.120.158101.

[16] P. L. Luisi, P. Walde, T. Oberholzer, Lipid vesicles as possible intermediates in the origin of life, Curr. Op. Coll. Int. Sci. 4 (1) (1999) 33–39. doi:10.1016/S1359-0294(99)00012-6.

[17] M. Kreysing, L. Keil, S. Lanzmich, D. Braun, Heat flux across an open pore enables the continuous replication and selection of oligonucleotides towards increasing length, Nature chemistry 7 (2015) 203–208. doi:10.1038/nchem.2155.

[18] P. Baaske, F. M. Weinert, S. Duhr, K. H. Lemke, M. J. Russell, D. Braun, Extreme accumulation of nucleotides in simulated hydrothermal pore systems, Proc. Natl. Acad. Sci. U.S.A. 104 (22) (2007) 9346–9351. doi:10.1073/pnas.0609592104.

[19] E. V. Koonin, W. Martin, On the origin of genomes and cells within inorganic compartments, Trends Genet. 21 (12) (2005) 647–654. doi:10.1016/j.tig.2005.09.006.

[20] B. Damer, D. Deamer, Coupled phases and combinatorial selection in fluctuating hydrothermal pools: A scenario to guide experimental approaches to the origin of cellular life, Life 5 (1) (2015) 872–887. doi:10.3390/life5010872.

[21] T. Furubayashi, N. Ichihashi, Sustainability of a compartmentalized host-parasite replicator system under periodic washout-mixing cycles, Life 8 (3) (2018) 10. doi:10.3390/life8010003.

[22] S. Matsumura, A. Kun, M. Ryckelynck, F. Coldren, A. Szilágyi, F. Jossinet, C. Rick, P. Nghe, E. Szathmáry, A. D. Griffiths, Transient compartmentalization of RNA replicators prevents extinction due to parasites, Science 354 (6317) (2016) 1293–1296. doi:10.1126/science. aag1582.

[23] J. S. Chuang, O. Rivoire, S. Leibler, Simpson’s paradox in a synthetic microbial system, Science 323 (5911) (2009) 272–275. doi:10.1126/science.1166739.

[24] A. S. Tupper, P. G. Higgs, Error tresholds for rna replication in the presence of both point mutations and premature termination errors, J. Theor. Biol. 428 (2017) 34–42. doi:10.1016/ j.jtbi.2017.05.037.

[25] Y. E. Kim, P. G. Higgs, Co-operation between polymerases and nucleotide synthethases in the rna world, PLoS computational biology 12 (11) (2016) e1005161. doi:10.1371/journal. pcbi.1005161.

[26] L. Geyrhofer, N. Brenner, Coexistence and cooperation in structured habitats, bioRxiv doi: 10.1101/429605.

[27] A. Zadorin, Y. Rondelez, Natural selection in compartmentalized environment with reshuffling, arXiv: 1707.07461.

[28] T. C. t. Michaels, A. J. Dear, T. P. j. Knowles, Stochastic calculus of protein filament formation under spatial confinement, New Journal of Physics 20 (5) (2018) 055007. doi: 10.1088/1367-2630/aac0bc.

[29] B. Houchmandzadeh, Giant fluctuations in logistic growth of two species competing for limited resources, Phys. Rev. E 98 (2018) 042118. doi:10.1103/PhysRevE.98.042118.

[30] D. L. Floyd, S. C. Harrison, A. M. van Oijen, Analysis of kinetic intermediates in single-particle dwell-time distributions, Biophys. J. 99 (2) (2010) 360–366. doi:10.1016/j.bpj.2010.04.049.

[31] A. Johnson-Buck, W. M. Shih, Single-molecule clocks controlled by serial chemical reactions, Nano Letters 17 (12) (2017) 7940–7944. doi:10.1021/acs.nanolett.7b04336.

[32] S. Karlin, H. M. Taylor, A first course in stochastic processes, Academic Press, 1975. doi: 10.1016/C2009-1-28569-8.

[33] B. J. Hindson, K. D. Ness, D. A. Masquelier, P. Belgrader, N. J. Heredia, A. J. Makarewicz, I. J. Bright, M. Y. Lucero, A. L. Hiddessen, T. C. Legler, et al., High-throughput droplet digital pcr system for absolute quantitation of dna copy number, Analytical chemistry 83 (22) (2011) 8604–8610. doi:10.1021/ac202028g.

[34] A. Dramé-Maigné, I. Golovkova, A. Zadorin, Y. Rondelez, Quantifying the performance of high-throughput directed evolution protocols, ArXiv: 1811.05288v1.

